# A maximum-type microbial differential abundance test with application to high-dimensional microbiome data analyses

**DOI:** 10.1101/2022.07.13.499972

**Authors:** Zhengbang Li, Xiaochen Yu, Hongping Guo, TingFang Lee, Jiyuan Hu

## Abstract

**Background:** High-throughput metagenomic sequencing technologies have shown prominent advantages over traditional pathogen detection methods, bringing great potential in clinical pathogen diagnosis and treatment of infection diseases. Yet, how to accurately detect the difference of microbiome profiles between treatment or disease conditions remains computationally challenging.

**Results:** In this study, we propose a novel test for identifying the difference between two high-dimensional microbiome abundance data matrices based on the centred log-ratio transformation of the microbiome compositions. The test p-value can be calculated directly with a closed-form solution from the derived asymptotic null distribution. We also investigate the asymptotic statistical power against sparse alternatives which are typically encountered in microbiome studies. The proposed **M**aximum-type test is **E**qual-**C**ovariance-**A**ssumption-**F**ree (**MECAF**), making it widely applicable to studies that compare microbiome compositions between conditions. Our simulation studies demonstrated that the proposed MECAF test achieves desirable power than competing methods while having the type I error rate well controlled under various scenarios. The usefulness of the proposed test is further illustrated with two real microbiome data analyses. The source code of the proposed method is freely available at https://github.com/JiyuanHu.

**Conclusions:** MECAF is a flexible differential abundance test and achieves statistical efficiency in analyzing high-throughput microbiome data. The proposed new method will allow us to efficiently discover shifts of microbiome abundances between disease and treatment conditions, broadening our understanding of the disease and ultimately improving clinical diagnosis and treatment.

## 1 Introduction

Human microbiota, a collection of microbes living on or inside human bodies, have shown to play fundamental role in human health and diseases, including diabetes, cancer, obesity, etc. (Turnbaugh et al. [2007], Ursell et al. [2012]). Two next-generation-sequencing (NGS) techniques, either 16S rRNA amplicon sequencing or shotgun metagenomic sequencing, are typically used to quantify the taxonomic and functional profiles and elucidate the association with diseases and phenotypes. Recently, the metagenomic NGS (mNGS) sequencing technique has been introduced to clinical diagnosis of infectious diseases (Gu et al. [2019], Dulanto Chiang and Dekker [2020], Govender et al. [2021]). It is emerging as a revolutionary technique to replace/supplement traditional culture-based and molecular microbiologic techniques that i) mNGS allows the parallel sequencing of hundreds of samples per run; ii) it provides an unbiased detection of bacteria, viruses, fungi, and parasites collectively; iii) This culture-free technology enables the identification of new species, etc.

In microbiome studies, it is of general research interest to study the microbiome profiles/features between different disease treatments or conditions. Various statistical methods have been proposed recently for examining the differential abundances (DA) (Cao et al. [2018], Banerjee et al. [2019], Zhao et al. [2018], Lin and Peddada [2020]). These methods can be categorized into univariate and multivariate approaches depending on whether microbial features are analyzed individually or in a set-based fashion. For example, Lin and Peddada [2020] proposed ANCOM-BC under a linear regression framework to conduct DA analysis for the assessed taxa individually. However, multiple comparison procedures need to be conducted afterwards for these univariate methods, which largely hinders the statistical power (Hu et al. [2018]). Alternatively, we can assess the microbial features as a set in order to enhance the statistical power. Typically, the microbial abundances are normalized towards the total counts to make the microbial proportions (or called relative abundances, RA) comparable between samples. The normalized data has summation of the features equal to one, termed compositional in microbiome studies (Mandal et al. [2015], Gloor et al. [2017]). Directly applying standard multivariate statistical methods developed for unconstrained data to compositional data may result in inappropriate or misleading inferences. Cao et al. [2018] proposed a two-sample test for assessing the difference between two high-dimensional microbial composition matrices and treat all microbial features (the microbiome profile) as a set. Banerjee et al. [2019] proposed an adaptive test for comparing microbiome compositions from two independent groups. Zhao et al. [2018] developed a generalized Hottelling’s test for paired microbiome composition data comparison. These methods can be applied to the full microbiome profiles and also microbial features which belong to the same upper-level taxonomic rank, gene family, or functional pathway. Yet, they either need strong assumption that the covariance matrices of compared compositions are equal Cao et al. [2018], or require time-consuming permutations to determine the statistical significance (Zhao et al. [2018], Banerjee et al. [2019]).

To address this challenge, we propose a two-sample **M**aximum-type **Equal**-**C**ovariance-**A**ssumption-**F**ree test named MECAF. This multivariate differential abundance test statistics relaxes the equal covariance assumption required by the test proposed in Cao et al. [2018]. The closed-form formula of the asymptotic null distribution largely resolves the computational burden in microbiome analysis. The method can be applied to analyze both taxonomic and functional profiles including microbial taxa (OTUs, strains, etc. from either shotgun metagenomic or 16S rRNA amplicon sequencing technique), functional pathways, and gene families. The performance of the proposed MECAF test is demonstrated through simulation studies and applications to the shotgun metagenome sequencing study of Clostridium difficile infection (CDI) (Vincent et al. [2016]), and the 16S rRNA amplicon murine microbiome study of type I diabetes (T1D) (Livanos et al. [2016]).

The rest of this article is as follows. In Section 2, we introduce the novel test statistics MECAF for conducting two-group comparison of microbiome compositions and summarize the superior asymptotic properties. In Section 3, we carry out extensive simulations to estimate the empirical type I error rate and statistical power for the proposed test in comparison with competing methods. Two real data applications are conducted in Section 4 and a concluding remark in given in the last section. All the theoretical derivations are detailed in the Online Supplemental Material.

## 2 Methods

### 2.1 Notation and specification of test hypothesis

In this article, we consider microbiome compositions from two independent groups. The notation used in this manuscript has been summarized in Table 1. Specifically, for subject *i* from group *g*(*g* = 1, 2), denote the *n*_*g*_ independently observed composition vectors as 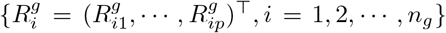 with length of *p*, and the *j*th component (taxon) of the vector 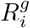 as 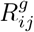, where 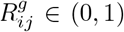. Of note, zero proportions are imputed by a pseudo positive proportion prior to conduct the analysis. Then the compositional constraints can be expressed as 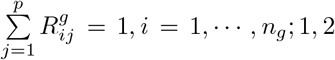. Obviously 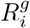 represents compositions lie in the *p* − 1 dimensional simplex 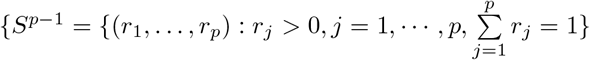 *R*^1^ and *R*^2^ are the observed data matrices of dimension *n*_1_ × *p* and *n*_2_ × *p* respectively from the two groups. Let 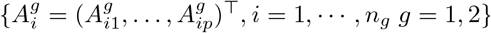 denote the *n*_*g*_ unobserved absolute abundances of the microbiome. The numerical relationship between the absolute abundance matrix and composition matrix is:

**Table 1:** Notation Summary.

| Notation | Description |
| --- | --- |
| $g$ | Group indicator, $g = 1, 2$ |
| $n_g$ | Sample size for group $g$ |
| $i$ | $i$th sample, $i = 1, 2, \dots, n_1$ for group 1, and $i = 1, 2, \dots, n_2$ for group 2 |
| $p$ | Number of taxa in the microbiome data matrix |
| $R^g$ | Observed microbial relative abundances (RA) for group $g$ with dimension $n_g \times p$ |
| $R_i^g$ | Observed RA for subject $i$ for group $g$ |
| $A^g$ | Unobserved microbial absolute abundances (AA) for group $g$ with dimension $n_g \times p$ |
| $L^g$ | Unobserved Log transformation of AA (log-AA) for group $g$ with dimension $n_g \times p$ |
| $L_i^g, \mu_L^g$, and $\Sigma_L^g$ | Unobserved log-AA for subject $i$ from group $g$,<br>and $L_i^g$ *i.i.d* from distribution with mean $\mu_L^g$ and covariance matrix $\Sigma_L^g$ |
| $X^g$ | Observed centered log ratio transformation of relative abundances of group $g$ with<br>dimension $n_g \times p$ |
| $X_i^g, \mu_X^g$, and $\Sigma_X^g$ | Observed CLR transformation of RA for subject $i$ from group $g$,<br>and $X_i^g$ *i.i.d* from distribution with mean $\mu_X^g$ and covariance matrix $\Sigma_X^g$ |

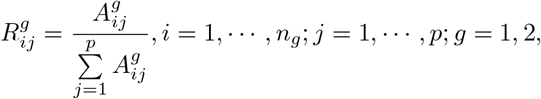

where 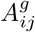 is the *j*th component of 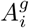. Suppose the log transformations of 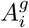, denoted by 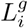, are i.i.d. from distributions with mean vectors 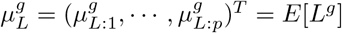 and covariance matrices 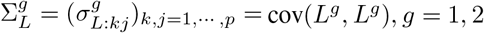. Cao et al. [2018] introduced a testable hypothesis to compare the log-mean absolute abundance vectors through the observed compositional data *R*^1^ and *R*^2^ by exploiting the centred log-ratio (CLR) transformation of the compositions. Denote the CLR transformation of 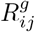 by

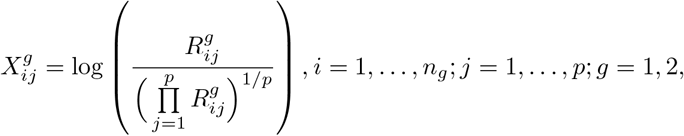

Assume that the CLR vectors 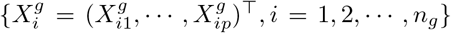 are i.i.d. from distributions with mean vectors 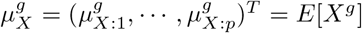, and covariance matrices 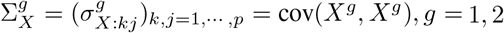. Then the testable hypothesis under the definition of compositional equivalence (please see Definition 1 in Cao et al. [2018]) is,

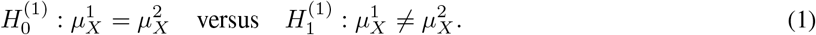

In this work, we consider another testable hypothesis of compositional equivalence as follows. It is straightforward that 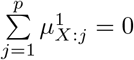, and 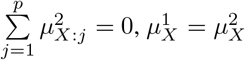 holds if and only is for *j* ∈ {1, …, *p* − 1}, as 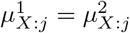. Therefore an equivalent hypothesis that only considers the first *p* − 1 components is,

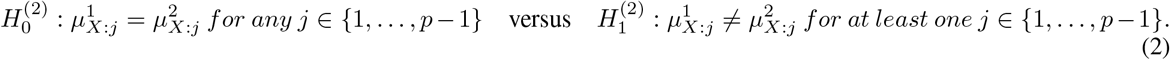

In the following, we will introduce our proposed test specifically for hypotheses (2). We also investigate the theoretical properties of the test statistics.

### 2.2 The proposed MECAF test

Cao et al. [2018] proposed one maximum-type two-sample test for high-dimensional compositions by assuming the covariance matrices of two groups being equal (see Equation-9 in Cao et al. [2018]). In practice, it is unable to assess the assumption if 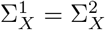 or not. Thus we consider a more general setting, where the equal covariance assumption is not required. For *j* ∈ {1, …,}th component(taxon), let 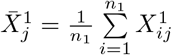, and 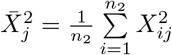 being the average of the CLR transformation of relative abundances. Our proposed test is given as

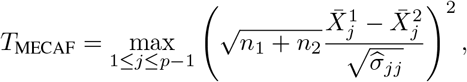

where 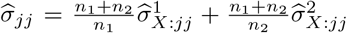, and 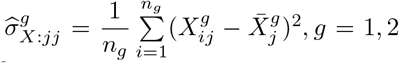 respectively. We name it the **m**aximum-type **e**qual-**c**ovariance-**a**ssumption-**f**ree (MECAF) test. As a maximum-type test statistics, it is in general better than sum-of-squares type statistics under sparse alternative. The assumption of the equal high dimensional covariance matrices for two groups are relaxed to allow for wider applicable conditions.

We successfully derived the asymptotic null distribution of *T*_MECAF_ given by

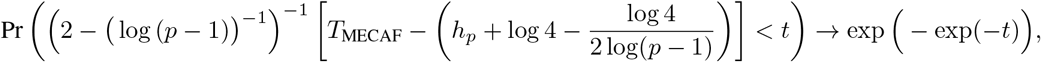

for any real number *t* as *n*_1_, *n*_2_, *p* → ∞, where 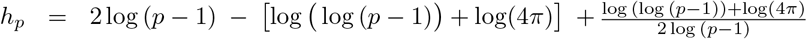. Denote *q*_*α*_ the (1 − *α*)-quantile of the derived distribution function exp (− exp(−*t*)). Namely, *q*_*α*_ = − log[log(1 − *α*)^−1^]. We can define an asymptotic *α*-level test denoted by

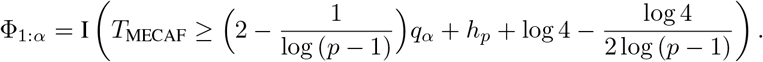

The null hypothesis 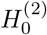 is rejected whenever Φ_1:*α*_ = 1. We also prove that the power of test Pr(Φ_1:*α*_ = 1) converges to 1 under some settings and 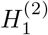 as *n*_1_, *n*_2_, *p* → ∞. All the detailed proof is given in the supplementary material.

## 3 Simulation Studies

### 3.1 Simulation setup

We conducted extensive simulations to evaluate the numerical performance of the proposed MECAF test comparing with competing methods under various scenarios. The simulation parameters were set up similarly as that in Cao et al. [2018] for the case of two independent samples in order to generate microbiome composition data. The log transformation of microbiome absolute abundance data *L*^1^ and *L*^2^ were first generated from the multivariate Gaussian distribution by assuming that 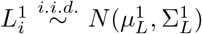 and 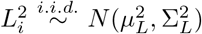. Then the raw absolute abundances *A*^1^, *A*^2^, relative abundances *R*^1^, *R*^2^, and CLR transformation of RA matrices *X*^1^, *X*^2^ can be generated accordingly with certain transformations detailed in Methods Section. We specify the location and covariance parameters for distributions 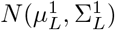 and 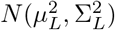 detailed as follows so that simulation data matrices can be generated with various covariance structures under the null and alternative hypotheses.

- **Specification of location parameters** 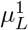 **and** 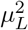. Following Cao et al. [2018], the components of 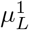 were drawn from the uniform distribution Uniform(0, 10). Each component of 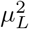 was set by 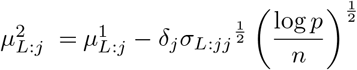. Here *δ*_*j*_ represents the signal, i.e., the difference in CLR means for component *j* between two groups. *s* = ⌞0*p*⌟, ⌞0.05*p*⌟, ⌞0.1*p*⌟, and ⌞0.2*p*⌟ components (taxa) and randomly chosen from *p* components to be the signal taxa and the corresponding *σ*_*j*_’s were randomly drawn from Uniform 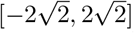. The other *σ*_*j*_’s were set as 0. We can see that *s* = ⌞0*p*⌟ corresponds to the null hypothesis setting and *s* = ⌞0.05*p*⌟, ⌞0.1*p*⌟, ⌞0.2*p*⌟ represent three alternative hypothesis settings. When *s* gets larger, there are more signal taxa in the microbiome compositions. *σ*_*L*:*jj*_ is the *j*’th diagonal component of the covariance matrix 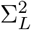 with specification as follows.
- **Specification of covariance matrices** 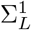 **and** 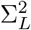. We included two types of covariance matrices, i.e., banded covariance matrix Σ_*B*_ and sparse covariance matrix Σ_*S*_ with the same parameters as those in Cao et al. [2018]. Three scenarios were considered to assess the impact of equal vs. unequal covariance matrices between two groups in the comparison of compositional mean vectors. Specifically, in Scenario 1, the covariance matrices of group 1 and 2 are set as 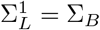, and 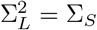 to represent the setting with unequal covariance matrices between groups; equal banded covariance matrices were considered in Scenario 2, i.e., 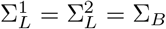 and equal sparse covariance matrices were considered in Scenario 3, i.e., 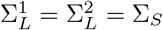.

### 3.2 Competing methods

By assuming that covariance matrices for the CLR of compositions being equal, i.e., 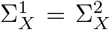, Cao et al.(2018) proposed a test for hypotheses (1) as 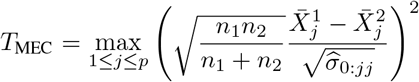, where 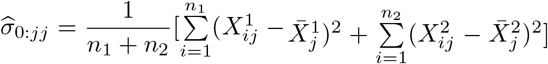. Since this is a **m**aximum-type test with **e**qual **c**ovariance assumption, we denote it the MEC test in this article. In addition, we also assessed the performance of the MEC statistics applied to the raw RA, the log transformation of RA, and the original AA. These three tests are obtained through replacing the CLR data used by MEC, i.e., *X*^*g*^, with *R*^*g*^, log(*R*^*g*^), and *A*^*g*^, *g* = 1, 2, denoting by MEC-Raw, MEC-Log, and MEC-Oracle respectively. MEC-Oracle is considered as the benchmark method in the simulation study (under equal covariance matrices assumption) as the true difference is simulated for the log-absolute abundances.

We set the sample size in the first group as *n*_1_ = 100, and increased the sample size in the second group *n*_2_ from 200 to 300. We increased the number of components (taxa) *p* 100 to 300 to demonstrate different relationship between *n* = *n*_1_ + *n*_2_ and *p*. We set the significance level as *α* = 0.05 in the simulation and 1000 replications were conducted to evaluate the empirical type I error rate and statistical power of the assessed methods under various settings.

### 3.3 Simulation results

Figure 1 shows the numerical performance of assessed methods under Scenario 1, where unequal covariance matrices are considered. All competing methods that require equal covariance matrix assumption, i.e., MEC-Oracle, MEC-Log, MEC-Raw, and MEC have inflated type I error rates. The type I error rate of MEC approaches 0.25 when *p* = 150, and *p* = 200. This indicates MEC-type tests are not applicable to data with unequal matrices. In comparison, the proposed MECAF test can control the empirical type I error rate around the nominal level of 0.05. The statistical power of MECAF increases with the proportion of signal taxa.

**Figure 1:**
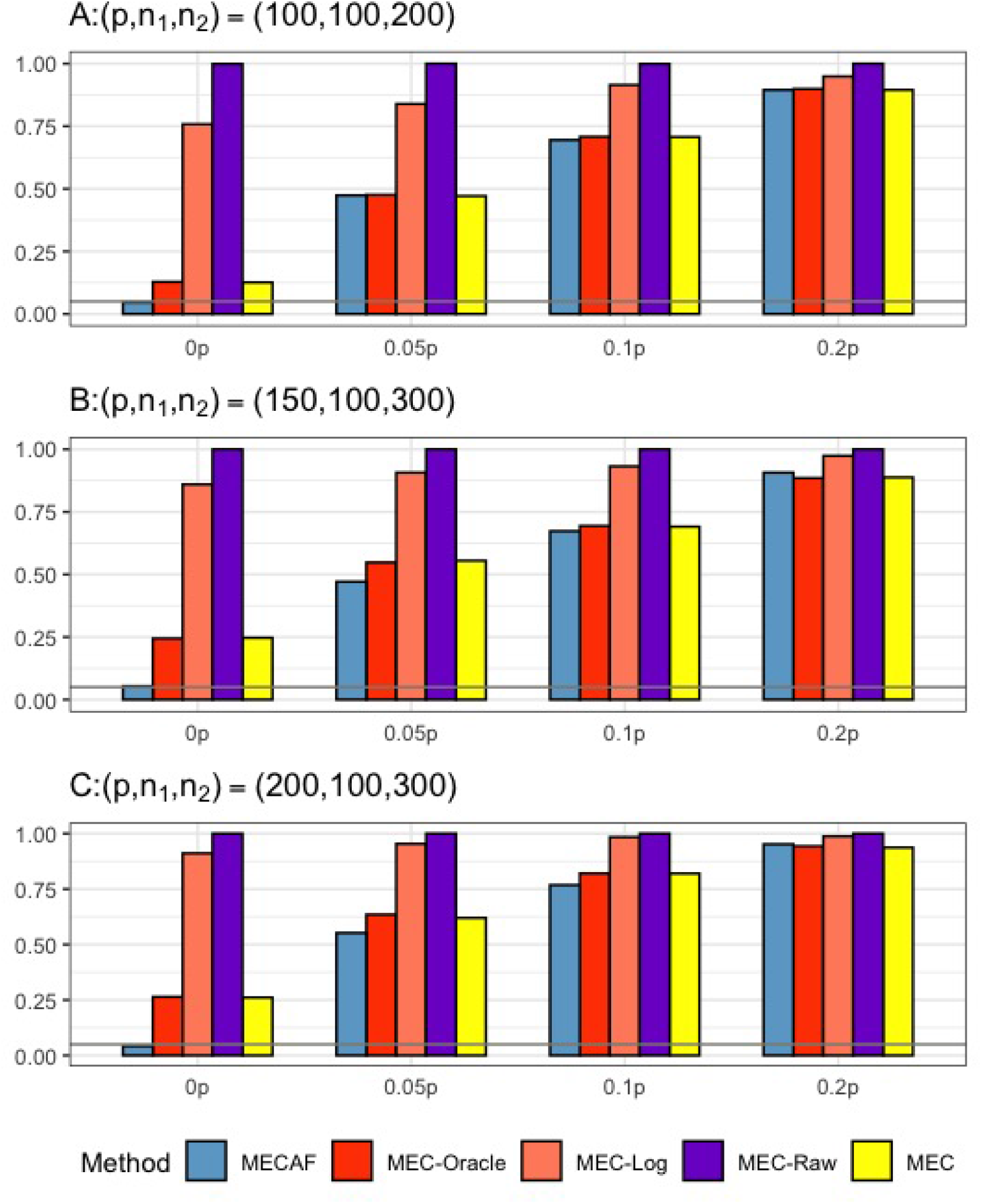
Simulation results for Scenario 1: unequal covariance matrices between compared groups. The empirical type I error rate (*H*_0_) and statistical power under three sparsity measures (*H*_*a*_ : 0.05*p*, 0.1*p*, and 0.2*p* respectively for MECAF and competing methods MEC-Oracle, MEC-Log, MEC-Raw, and MEC. A horizontal line with *α* = 0.05 was used to indicate the significance level. The number of taxa and sample sizes were set as follows: (A) (*p, n*_1_, *n*_2_) = (100, 100, 200); (B) (*p, n*_1_, *n*_2_) = (150, 100, 300); (C) (*p, n*_1_, *n*_2_) = (200, 100, 300).

Simulation results for equal banded covariance and equal sparse covariance scenarios are depicted in Figure 2 and Figure 3 respectively. As expected, all assessed methods have well controlled type I error rate under these simulation settings. MECAF and MEC achieved statistical power comparable to that of MEC-Oracle, with sparsity measure *s* ranging from ⌞0.05*p*⌟ to ⌞0.2*p*⌟. This indicates the statistical efficiency of the MECAF test. In comparison, MEC-Log and MEC-Raw have evidently smaller power than MEC-Oracle for all settings of two scnarios.

**Figure 2:**
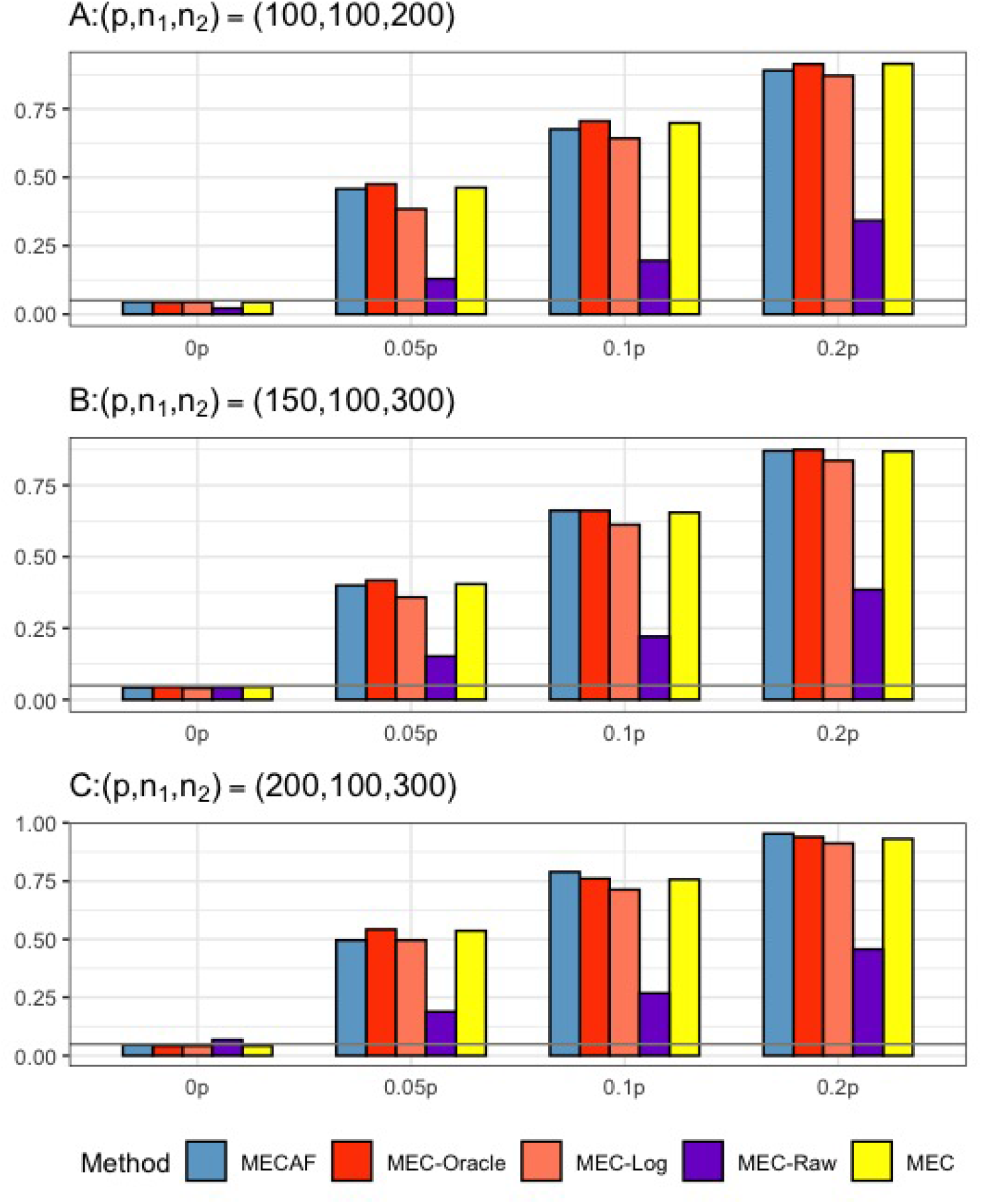
Simulation results for Scenario 2: same banded covariance matrix between compared groups. The empirical type I error rate (*H*_0_) and statistical power under three sparsity measures (*H*_*a*_ : 0.05*p*, 0.1*p*, and 0.2*p* respectively for MECAF and competing methods MEC-Oracle, MEC-Log, MEC-Raw, and MEC. A horizontal line with *α* = 0.05 was used to indicate the significance level. The number of taxa and sample sizes were set as follows: (A) (*p, n*_1_, *n*_2_) = (100, 100, 200); (B) (*p, n*_1_, *n*_2_) = (150, 100, 300); and (C) (*p, n*_1_, *n*_2_) = (200, 100, 300).

**Figure 3:**
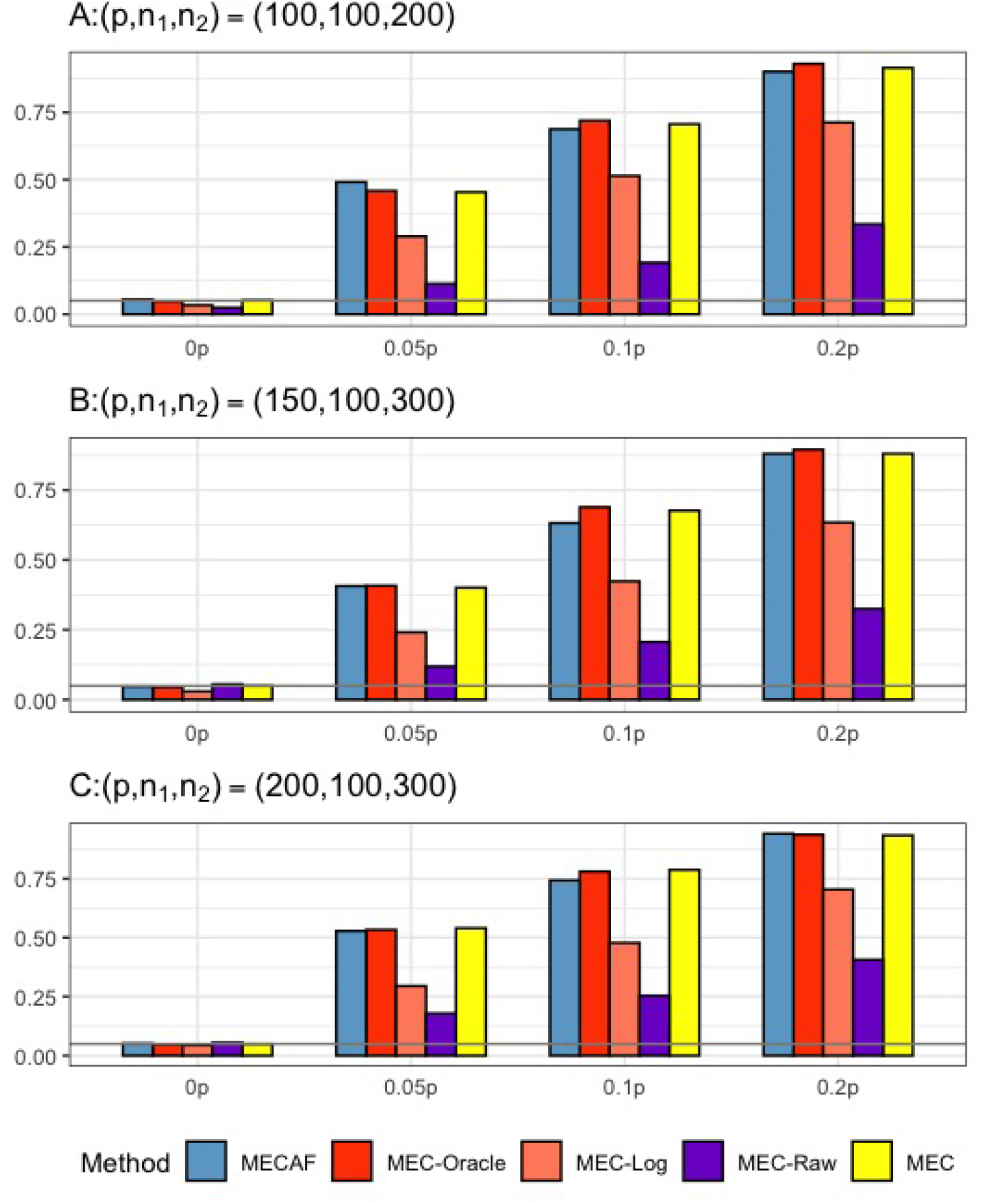
Simulation results for Scenario 3: same sparse covariance matrix between compared groups. The empirical type I error rate (*H*_0_) and statistical power under three sparsity measures (*H*_*a*_ : 0.05*p*, 0.1*p*, and 0.2*p* respectively for MECAF and competing methods MEC-Oracle, MEC-Log, MEC-Raw, and MEC. A horizontal line with *α* = 0.05 was used to indicate the significance level. The number of taxa and sample sizes were set as follows: (A) (*p, n*_1_, *n*_2_) = (100, 100, 200); (B) (*p, n*_1_, *n*_2_) = (150, 100, 300); and (C) (*p, n*_1_, *n*_2_) = (200, 100, 300).

In summary, MECAF has a well controlled type I error rate for two group comparison of mean composition vectors either with equal or unequal covariance matrices. The statistical power is desirable under all scenarios with various sparsity measures.

## 4 Applications to two microbiome studies

Here, we first apply the proposed MECAF test to the shotgun metagenomic sequencing data from the *Clostridium difficile* infection (CDI) study (Vincent et al. [2016]). Since the test is also applicable to microbiome abundance data generated from the 16S rRNA amplicon sequencing technology, we further illustrate our proposed method in a murine microbiome study of type I diabetes (T1D) (Livanos et al. [2016]).

### 4.1 Analysis of the CDI metagenomic dataset

Vincent et al. [2016] conducted a prospective study to investigate the intestinal microbiota dynamics over time among 98 hospitalized patients at risk for CDI, a leading infectious cause of nosocomial diarrhea. Patients were followed up to 60 days and a total of *N* = 229 fecal samples (averaging 2.34 samples per subject) were examined by the shotgun metagenomics sequencing platform. The bioinformatics pre-processing steps were detailed at Vincent et al. [2016], and the processed microbial counts and meta data are available in the R package “curatedMetagenomicsData”(Version 1.16.1) from the Bioconductor through running the function *curatedMetagenomic-Data(‘VincentC 2016*.*metaphlan bugs list*.*stool’,dryrun = FALSE)* (Pasolli et al. [2017]). Zero counts were imputed with 0.5 before converting counts to relative abundances for taxa from taxonomic ranks of phylum, class, order, family, genus, and species (strains) respectively. In this secondary data analysis, we aim to examine whether there are shifts in the microbial relative abundances between patients with CDI or asymptomatic *C. difficile* colonization (CDI group, N = 8 subjects) and patients without (control group, N = 90 subjects) i) upon hospitalized (baseline), ii) at one week of hospitalization, and iii) over one week of hospitalization respectively. The latest sample at each time window was included for each patient.

Figure 4 shows the available samples at each assessed time window (Fig. 4 A), number of taxa observed at each taxonomic rank (Fig. 4 B), the differential abundance test results of MECAF test and competing methods (Fig. 4 C-E). P-value of *<* 0.05 was indicated as statistical significance. Of note, MEC-Oracle was not included into comparison since the absolute abundance data required by MEC-Oracle are unobservable in real data. The results of MECAF indicate significant difference in the microbiome compositions between CDI and control patients at baseline and week one. The significance is consistent for most of the taxonomic ranks, and stronger signal is depicted at the lower ranks. The difference, though seems disappear after one week of hospitalization, where only the test at the species(strain) level is significant. In comparison, MEC does not detect significant difference in microbiome compositions except at the species and strain level at baseline and over one week, with less stringent p-values reported. At week one, MEC detects microbial composition difference at the family, genus and species (strain) level, but not at the phylum, class, or order level. MEC-Raw has similar results as MEC, and MEC-log reported similar results as those of MECAF with larger p-values. In summary, we observe more consistent findings from the MECAF test over six taxonomic ranks. The corresponding p-values are in general smaller than competing methods, indicating statistical efficiency gain over other methods.

**Figure 4:**
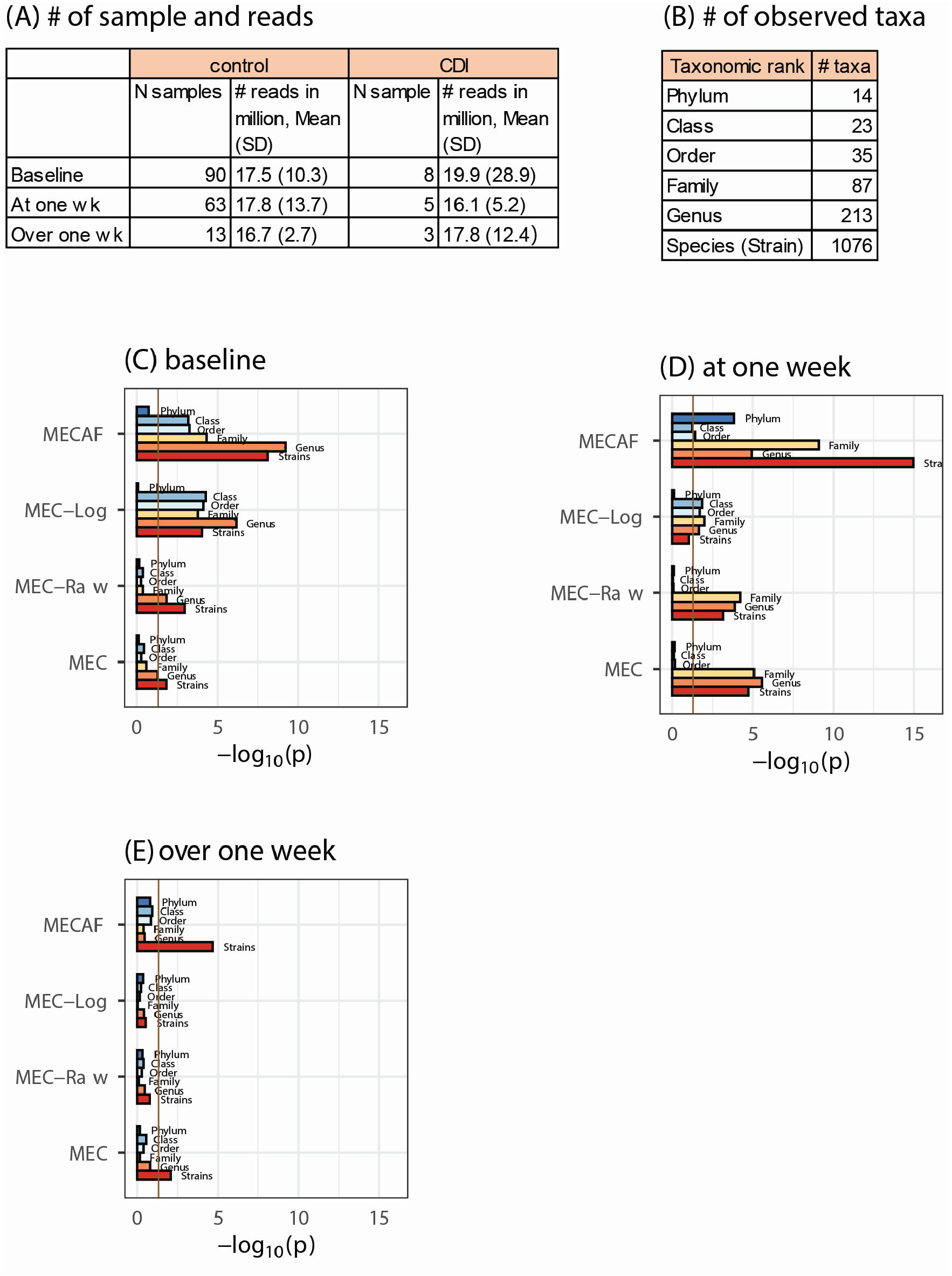
DA analysis of the CDI metagenomic dataset. (A) The number of samples and microbial reads were summarized; (B) Number of observed taxa at each of the taxonomic rank; (C-E) P-values of DA test with MECAF and competing methods at baseline, one week, and over one week, respectively. P-values were −log_10_ transformed to better illustrate the statistical significance where the vertical line of − log_10_ 0.05 indicated a p-value equal to 0.05.

### 4.2 Analysis of the MICE 16S rRNA amplicon microbiome data

Livanos et al. [2016] carried out a murine microbiome study to investigate the effect of early-life antibiotic exposure on the alteration of gut microbiota composition. Here, we re-examined the 16S microbiome abundance profile from the early-life sub-therapeutic antibiotic treatment (STAT) group and the control group that received no antibiotic exposure. The abundances were compared between the two groups at each of the four assessed time points, i.e., week 3, 6, 10, and 13 respectively for female and male mice using MECAF test and competing methods.

The available samples and number of taxa observed were shown in Fig. 5 A-B, which illustrate the circumstance of *n < p* (number of samples *<* number of taxa) most often encountered in microbiome data analysis. The differential abundance analysis results from the phylum to genus rank are depicted in Fig. 5 C and Fig. 6 for female and male mice respectively. The result from MECAF indicated that in female mice, the abundance profile is significantly different in STAT group from the control group from week 3 to 6 for almost all taxonomic ranks. The significance is weaker at week 10 and 13, indicating the recovery of gut microbiota in the STAT mice upon maturation. In comparison, the significant difference is detected by MECAF in male mice over the four assessed time points. This result is consistent with Livanos et al. [2016] in which the alpha and beta diversity measures were compared between groups over time. MEC has similar results with that of MECAF, yet with slightly larger p-values. MEC-Log did not detect significance in female mice from week 6-10 for most of the taxonomic ranks. MEC-Raw, either did not detect significant difference (female mice from weeks 3-10), or reported weaker signals (male mice, weeks 3, 10, 13).

**Figure 5:**
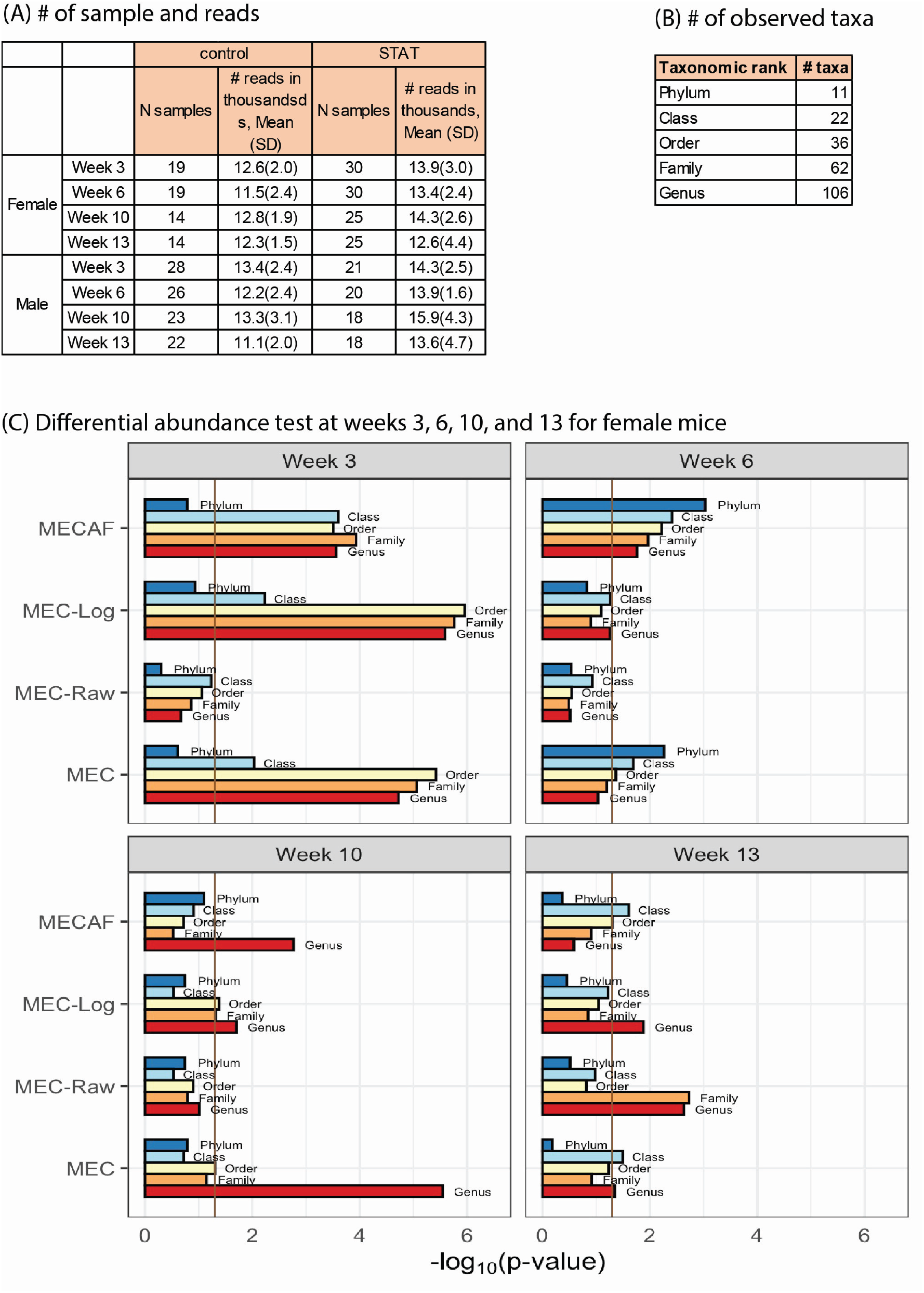
DA analysis in female mice of the murine microbiome study. (A) The number of samples and microbial reads were summarized seperately for female and male mice; (B) Number of observed taxa at each of the taxonomic rank; (C) P-values of DA test with MECAF and competing methods at four assessed time points. P-values were −log_10_ transformed to better illustrate the statistical significance where the vertical line of −log_10_ 0.05 indicated the significance level.

**Figure 6:**
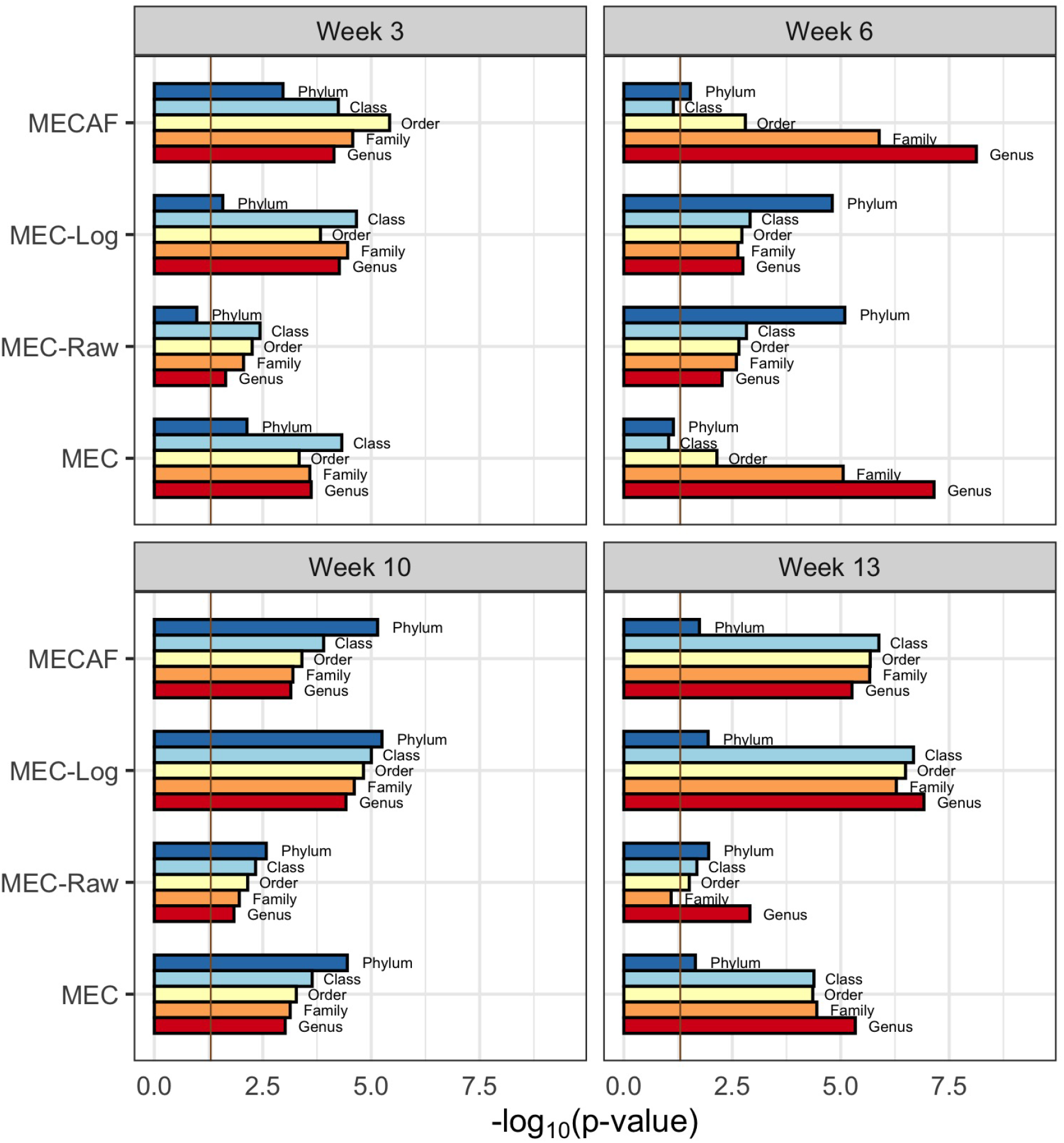
DA analysis in male mice of the murine microbiome study. P-values of DA test with MECAF and competing methods at four assessed time points were shown. P-values were −log_10_ transformed to better illustrate the statistical significance where the vertical line of − log_10_ 0.05 indicated the significance level.

## 5 Discussion

In this article, we propose a novel test named MECAF for the two-sample test of high-dimensional compositions. The test statistics is developed based on the centred log-ratio transformation of the compositions following Aitchison [1982] and Cao et al. [2018]. The asymptotic null distribution of the test statistic is derived and the power against sparse alternatives is investigated. The derived null distribution allows for the closed-form solution of statistical significance and largely resolves computational burden. Simulation results show that the proposed method is evidently more powerful than competing methods when the covariance matrices differ between groups, and comparable performance is achieved when the groups have equal covariance. Two real data applications have illustrated the usefulness of the proposed method.

The MECAF test extends MEC proposed by Cao et al. [2018] by relaxing the assumption of equal covariance matrix structure between groups. Therefore, MECAF can be applied to a wider set of circumstances. In the real data applications, we applied MECAF to compare the microbiome abundances aggregated to each taxonomic rank. In practice, we can also apply MECAF to assess the composition for a give sub-tree or a subset of the microbiome taxa (Shi and Li [2017]). As a future direction, we will aim to extend MECAF test to accommodate repeated measures from each individual for group comparisons.

## Author Contributions

ZL developed the proposed method, performed theoretical proof and simulation studies, and manuscript writing. XY performed simulation and real data analyses and manuscript writing. HG performed theoretical proof and manuscript writing. TL performed simulation analyses. JH conceptualized the ideas for the proposed method, simulations, real data analyses and manuscript writing. All authors read and approved the final manuscript.

## Acknowledgments

The authors would like to thank the anonymous reviewers and editors for the valuable comments and suggestions.

## Supplemental Data

Please check the online supplementary materials for the derivation of the asymptotic null distribution of the MECAF test statistics, and the asymptotic statistical power under sparse alternatives.

## Data Availability Statement

The metagenomics abundance data of the CDI study is readily available through the R package “curatedMetagenomicsData”(Version 1.16.1) from the Bioconductor (https://bioconductor.org/packages/release/data/experiment/html/curatedMetagenomicData.html). The 16S rRNA amplicon sequencing data from the murine T1D study is publicly available at EBI with accession number ERP016357.

